# 3D Printed Multifunctional Bioadhesive Patch with Intrinsic Bioelectronic Properties for Decoding Electromechanical and Anisotropic Cardiac Microenvironment

**DOI:** 10.1101/2024.07.05.601338

**Authors:** Sayan Deb Dutta, Tejal V. Patil, Ki-Taek Lim

## Abstract

Fabricating anisotropic multifunctional bioadhesive patches with tunable mechanical stiffness, electrical conductivity, antimicrobial activity, and modulating cellular behavior is crucial for the successful management of cardiac tissue injury and boosting immunogenic microenvironments. Direct ink writing (DIW)-based 3D printing holds tremendous potential for developing electroactive cardiac patches (ECPs) with anisotropic microarchitecture. Inspired by the native myocardium, we developed a multifunctional and anisotropic ECP with tunable stiffness by incorporating a highly conductive graphene oxide/nanodiamond (GO@ND) complex into a biocompatible carboxymethyl chitosan/polyvinyl alcohol (CSA) matrix for regulating immunogenic and cardiomyogenic cues. The incorporation of GO@ND enhanced the electrical conductivity (∼22.6 S mm^-1^) with high interfacial toughness (>250 MJ m^-1^) and improved the printability (*n* = 0.5) with concentration-dependent self-assembly into the CSA matrix. We observed that electrical stimulation (EFs; 250 mV/20 min/day) through nanoengineered CSA resulted in broad-spectrum antibacterial activity against *E. coli* and *S. aureus* by 99.29% and 98.74%, respectively, via sustained release of curcumin (Cur). Moreover, the electromechanical study revealed that CSA with higher stiffness (∼6.2 kPa) activated cytoplasmic YAPs during macrophage polarization. Besides, stiffness and EFs regulated human cardiomyocyte differentiation through anisotropic force-driven early activation of Vinculin, triggering the phosphorylation of NFATc3 and activating Lamin A/C in a YAP-dependent manner. Based on these findings, we anticipated that the fabricated nanoengineered patch had tremendous potential for regulating the electro-cardiomyogenic microenvironment with multifunctional abilities.

## INTRODUCTION

Anisotropic cardiac patches with intrinsic bioelectronic properties display a promising strategy for treating several heart diseases, including myocardial infarction (MI), cardiac arrhythmia (CA), and ventricular tachycardias (VTs). Also, smart cardiac bioelectronics is beneficial for monitoring the heartbeat, and anisotropic scaffolds mimic the ECM functions of the native myocardium. ^1–3^ In this context, conductive nanomaterial-based smart hydrogels could be used to enhance the conductivity and sensitivity of cardiac patches. Conductive nanomaterials, such as graphene, carbon nanotubes (CNT), titania carbide (MXene), and silver nanowires (AgNWs), have been used in combination with conductive biopolymers to restore damaged myocardium and epicardium. Doping of conductive materials with other nanomaterials improved the conductivity by adding or removing electrons, thereby maintaining conductivity.

Previous reports suggest that biophysical stimulation alters macrophage metabolism and induces polarization toward different subtypes. ^4^ For instance, macrophage fate can be directly regulated through changes in voltage-sensitive ion channels and receptors with appropriate biophysical or biochemical signals using conductive biomaterials. ^5–7^ Furthermore, surface charge and biomaterial topography have a positive role in macrophage polarization by eliciting specific immune responses. ^8^ Although several advancements have been devoted to controlling the pro-inflammatory or anti-inflammatory activation of macrophages, the exact role of electro-mechanotherapy in regulating macrophage polarization and subsequent tissue regeneration has yet to be explored. In this study, we examined the combinational effect of external EF and electroactive scaffold anisotropy on macrophage polarization toward cardiac tissue regeneration.

Hydrogel’s viscoelasticity and controllable stiffness play important roles in cell adhesion, migration, proliferation, and differentiation. ^9, 10^ Chemical crosslinking and a tougher hydrogel network also facilitate dynamic bond formation, which contributes to the self-healing and elasticity of the hydrogel matrix suitable for regulating cell fate.^10, 11^ To date, various ECM-mimicking hydrogel platforms with tunable stiffness have been demonstrated to regulate cardiomyocyte (CM) fate via altering metabolism or up-or down-regulating myogenesis-specific transcription factors. For instance, Young and Engler^12^ reported that hyaluronic acid (HA) hydrogels with a stiffness of ∼10 kPa displayed early activation of NKX2.5 while downregulating the expression of cTnT. In another study, polyacrylamide (PA) hydrogel with a stiffness ranging from 2-33 kPa exhibited no noticeable change in a-actinin expression. Still, higher PA stiffness (>10 kPa) reduced the sarcomere length significantly in neonatal rat cardiomyocytes (NRCMs). ^13^ Recently, Finklea et al. reported that a PEG-fibrinogen microsphere with extremely low stiffness (<250 Pa) promoted differentiation of human induced pluripotent stem cell (hiPSC)-derived cardiomyocytes (hiPSC-CMs). ^14^ Despite remarkable advancements in cardiovascular research, the role of electro-mechanostimulation, i.e., the synergistic role of electrical stimulation with mechanical (=hydrogel stiffness and topography) stimulation in hCMs proliferation and differentiation, is ill-explored.

In this study, we fabricated a nanoengineered conductive hydrogel, and we observed that nanofiller addition positively regulated the hydrogel’s conductivity, anisotropy, and stiffness, which would be a promising approach for studying the hCM’s fate. To enhance the conductivity, we doped graphene oxide (GO) with nanodiamonds (ND). The fabricated GO@ND complex displayed higher electrical conductivity when combined with CSA hydrogel. The as-fabricated anisotropic hydrogel patch showed superior mechanical and adhesive properties owing to the addition of GO@ND with varying stiffness. The hydrogel patch was found to be biocompatible with human ventricular cardiomyocytes (hCMs). Electromechanical stimulation through the nanoengineered hydrogel revealed that hCM differentiation is mainly regulated by scaffold anisotropy and stiffness, which are responsible for cellular mechanosensing.

## RESULTS AND DISCUSSION

### Fabrication of anisotropic patch with intrinsic bioelectronic properties

In this study, we developed a nanocomposite of GO@ND and used it as a reinforcing agent for CSA hydrogel to improve conductivity, sensitivity, and biocompatibility. **Fig. 1(a)** schematically shows the synthesis of GO@ND nanocomposite and anisotropic hydrogel patch fabrication. The morphology of the GO@ND nanocomposite was analyzed through transmission electron microscopy (TEM), field-emission scanning electron microscopy (FE-SEM), and atomic force microscopy (AFM). As shown in **Fig. 1b(i)**, the ND particles were assembled onto the thin sheets of GO. The ND particles form clusters on the GO surface, which was further confirmed by observing the FE-SEM and AFM morphologies **(Fig. 1b (ii, iii))**. The assembly of ND particles on the GO nanosheet will contribute to the enhancement of ‘metal-like’ electrical conductivity.

**Figure 1.**
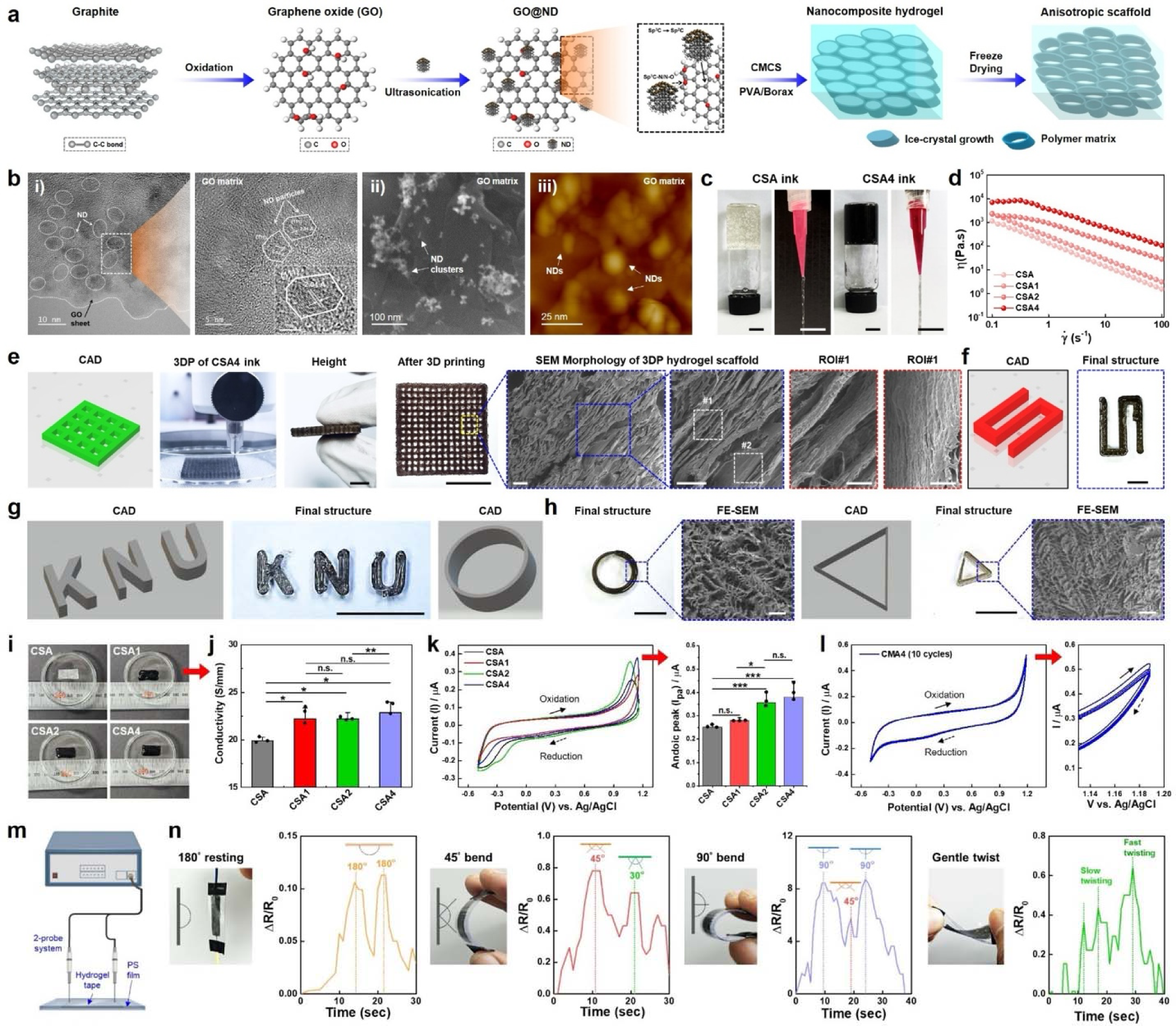
Characterization of the GO@ND and its nanocomposite hydrogels. **(a)** Schematic illustration of the GO@ND nanocomposite preparation. **(b)** HR-TEM, SEM, and AFM images of the GO@ND nanocomposites. Scale bar: 5, 10, 25, and 100 nm. **(c)** Demonstration of extrudability of the CSA (control) and CSA4 (experimental) inks for 3D printing. Scale bar: 5 and 10 mm. **(d)** Representative viscosity *vs.* shear rate of the fabricated hydrogels at RT showing the shear-thinning behavior. **(e)** Comparison of 3D printing performance of the CSA4 hydrogel ink with CAD model. Structures were printed and freeze-dried, which shows the aligned surface morphology. Scale bar: 10 mm, 100, and 10 µm. **(f, g)** Demonstration of 3D printing of the ‘*S*’ alphabet and ‘letter’ using CSA4 hydrogel ink. **(h)** 3D printing process and FE-SEM images of the tubular and trigonal geometry showing the lamellar structure with anisotropic pores. Scale bar: 10 mm. **(i, j)** Conductivity test of the developed hydrogel inks. Digital photographs of the hydrogels used in conductivity study. **(k, l)** Cyclic voltammetry test of the fabricated hydrogels showing the increased anodic current (I*_pa_*) flow. Data represented as mean ± s.d. of triplicate (*n* = 3) experiments, statistical significance considered at \**p* < 0.05, \*\*\**p* < 0.001 with One-way ANOVA test. **(m, n)** The sensing performance of the hydrogel. The sensing device was fabricated by taking the hydrogel tape (dimension: 20 × 5 × 2 mm^3^), which was enclosed by a thin layer of polystyrene (PS, thickness: 0.15 mm). The relative change in resistance after bending at various angles (30°, 45°, and 90°) and twisting.

The CSA hydrogel was fabricated with varying concentrations (0 to 4 wt.%) of GO@ND after confirming the electrical conductivity. The CMCS was crosslinked with PVA via borax, and the GO@ND was physically crosslinked with PVA and CMCS through inter- and intramolecular hydrogen bonding. The hydrogels were cast in cylindrical molds to form a thin film (∼0.5-1 mm), while the physically crosslinked hydrogels were loaded onto the printing cartridges **(Fig. 1(c))**. The shear-thinning behavior of the nanoengineered hydrogels (CSA to CSA4) was assessed through a flow curve (viscosity vs. shear rate) at room temperature. We observed that all the fabricated hydrogels exhibited shear-dependent thinning behavior within a measurement range of 0.1 to 100 s^-1^ **(Fig. 1(d))**. This is due to the ability of the GO@ND to contribute to the overall viscoelasticity, ^15^ suggesting that the hydrogels were printable.

Next, we examine the 3D printing performance of the fabricated hydrogel inks. We have shown the printing ability of the CSA4 (CSA with 4 wt.% GO@ND) as it showed good printing properties among all the composites. As shown in **Fig. 1(e)**, the CSA4 hydrogel showed good printing, and after freeze-drying, the 3D-printed scaffolds displayed lamellar pore geometry aligned with the printing direction. This phenomenon can be explained by the fact that carbon and non-carbon-based colloidal 2D nanomaterials have been shown to exhibit shear-induced self-assembly in the polymer matrix, resulting in aligned surface topography with tunable stiffness. ^15–17^ We also observed that the microporous lamellar structure was further confined into fiber-like morphology under high-resolution SEM, confirming the formation of aligned topography. We also tested various geometries, such as ‘S’-shaped and letter-based (KNU) geometry, to examine the printability of the CSA4 hydrogel ink **(Fig. 1(f, g))**. Similar to the square geometry (10 x 10 x 5 mm^3^; 30% infill density), the tubular (10 x 5 mm^2^; 0% infill density) and triangular (10 x 10 x 10 mm^3^; 0% infill density) geometries also displayed anisotropic scaffold morphology upon freeze-drying, which was probably due to the GO@ND assembly in the printing process **(Fig. 1(h))**. These results suggested that the CSA/GO@ND hydrogels are printable and exhibit an anisotropic surface, which would be beneficial for the growth and development of cardiomyocyte cells. ^18, 19^

The electrical properties of conductive biomaterials are essential for maintaining cell proliferation, migration, and differentiation, especially for cardiomyocytes. ^20, 21^ To access the conductive properties of the fabricated nanocomposite hydrogels, we used the 4-probe station and cyclic voltammetry methods. The test hydrogel samples (20 x 10 x 5 mm^3^) were photographed and shown in **Fig. 1(i)**. Notably, all the hydrogels containing GO@ND (1, 2, and 4 wt.%) exhibited better electrical conductivity (*p < 0.05) than the control hydrogel (CSA; no GO@ND). We found that the CSA4 hydrogel (∼22.61 S mm^-1^) showed significantly higher (*p < 0.05) conductivity than the CSA (∼19.47 S mm^-1^), suggesting the positive role of GO@ND in conductivity **(Fig. 1(j))**. To confirm the conductivity, we then studied the electrochemical performance of the fabricated hydrogel films, and the result is shown in **Fig. 1(k)**. Interestingly, with the increasing GO@ND content, the hydrogels displayed an enhancement of the peak anodic current (high in the case of CSA4, ***p < 0.001) at a scan rate of 50 mV s^-1^, suggesting a greater and undisrupted electron flow. ^22, 23^ Moreover, the outstanding conductivity of CSA4 was also reflected in cyclic current stability **(Fig. 1(k))**, which was due to the homogeneous distribution of GO@ND in the CSA matrix. ^24^

The electroactive patches can be used to monitor pressure, motion, and even deformation (=bending) sensitive events, which has a significant impact on biomedical electronics. ^25, 26^ However, most hydrogels display poor deformation-induced sensing performances toward cardiac bioelectronics development. Herein, we developed a highly conductive GO@ND-decorated 3D printable hydrogel ink for precise sensing of deformation/bending, one of the mimicking features of cardiac physiology. A schematic illustration of the sensing experiment is shown in **Fig. 1(m)**. We observed that by increasing the bending angle from 0° to 90°, a fast change in resistance (ΔR/R_O_) was observed. In particular, we were able to detect the change from 0° to 45° to 90° by monitoring the small and large peaks, respectively **(Fig. 1(n))**. A similar observation was noted for gentle twisting experiments, suggesting the excellent sensing performance of the fabricated hydrogels.

### Mechanical and adhesive properties of the fabricated hydrogels

Tensile measurements were conducted to examine the mechanical properties of the developed hydrogel films. As illustrated in **Fig. 2(a)**, the tensile stress gradually increased as we went from CSA to CSA4 (with increasing GO@ND content) samples. The CSA4 hydrogel displayed significantly higher tensile strength (∼29.1 MPa; ***p < 0.001) and elastic modulus (∼16.2 MPa; ***p < 0.001) than other hydrogel films **(Fig. 2(b))**. The superior tensile property of the CSA/GO@ND films was due to the homogenous distribution of GO@ND in the CSA matrix and its ability to form inner-connecting networks of bulk polymers, resulting in good self-healing and stretchability.

**Figure 2.**
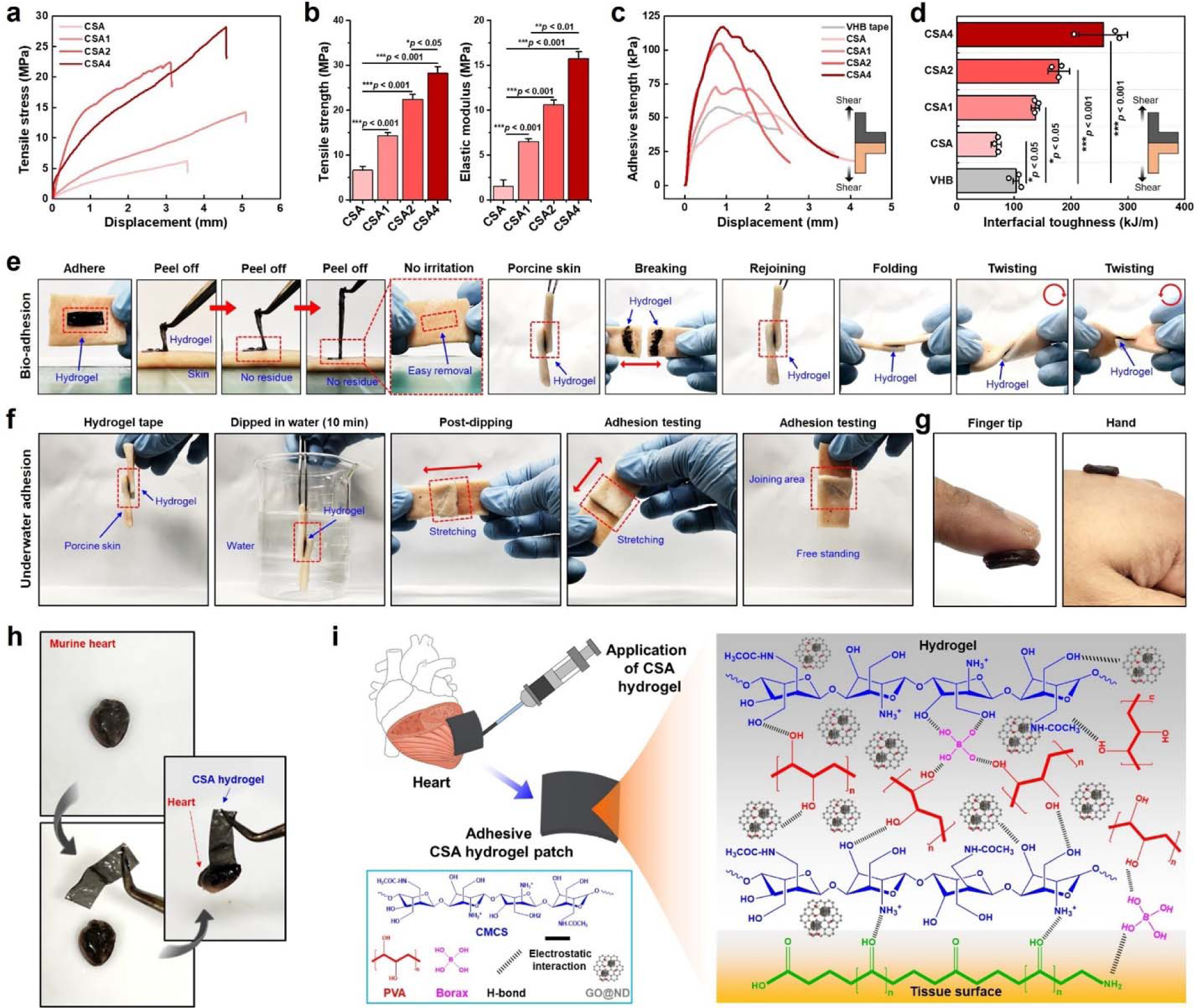
Mechanical and adhesive behavior of the CSA hydrogel inks. **(a, b)** Tensile stress-strain curve, ultimate tensile strength (UTS), and elastic modulus of the hydrogel films. **(c)** Interfacial adhesive test and **(d)** toughness of the fabricated hydrogels showing excellent adhesive strength. 150 mg hydrogel sample was tested for each group. Data represented as mean ± s.d. of triplicate (*n* = 3) experiments, statistical significance considered at \*\*\**p* < 0.001 with One-way ANOVA test. **(e)** Demonstration of bio-adhesion of the CSA4 hydrogel tape using the porcine skin sample (i). The CSA4 hydrogel was able to stick to the cut surfaces of the porcine skin (ii) even after bending or twisting. **(f)** Demonstration of underwater adhesion behavior of the CSA4 hydrogel tape. **(g)** Digital photographs of the CSA4 hydrogel adhesion on human skin surface. **(h)** Adhesive test on murine cardiac tissue. **(i)** Schematic illustration of the adhesive mechanism of CSA/GO@ND hydrogel tape.

The adhesive behavior of the fabricated hydrogel film was tested using multiple adhesive tests, such as the lap-shear test, the interfacial adhesive test, and the underwater bioadhesive test. The interfacial adhesive test was conducted to examine the tissue-intermediate adhesiveness, often useful for sealing internal organs. As shown in **Fig. 2(c)**, the adhesive strength increased with increasing GO@ND content and keeping the hydrogel films at physiological temperature (∼37°). The interfacial toughness of the CSA4 hydrogel (>250 MJ m^-1^, ***p < 0.001) was found to be the highest among all hydrogels and also greater than commercial VHB tape (∼100 MJ m^-1^) **(Fig. 2(d))**.

To verify the adhesiveness, we further performed the adhesive test using porcine skin. Interestingly, the CSA4 hydrogel was quickly (<10 s) adhered to the skin surface, as verified by the down-pulling method. Also, the hydrogel film was gently detached from the skin surface without any residual trace **(Fig. 2(e))**. Moreover, the hydrogel films, when inserted between overlapping skin samples, are tightly adhered to even after moving with force. Once the skin samples were forcefully detached, the residual hydrogel was found stuck on both ends of the skin samples, conferring extreme adhesion. Furthermore, the skin samples, when rejoined and twisted in a clockwise and anticlockwise manner, did not slip out, suggesting the self-healing ability of the hydrogels. We also tested the hydrogel’s adhesiveness by dipping it in tap water for 10 min **(Fig. 2(f))**, which showed non-degradable behavior and could be useful for sealing wet tissue, such as a punctured heart or coronary arteries. Finally, as a proof-of-concept, we tested the adhesive performance of the CSA4 hydrogel tap using human fingers and hands, as demonstrated in **Fig. 2(g)**. Digital photographs of the murine heart adhesive test are illustrated in **Fig. 2(h)**. Taken together, we anticipated that GO@ND-reinforced CSA hydrogels hold tremendous potential as tissue adhesives and could be beneficial for treating cardiac injuries. **Fig. 2(i)** schematic represents the bioadhesion mechanism of GO@ND/CSA hydrogel tape.

### Electrical stimulation enhanced the antibacterial effect of endocarditis

Infective endocarditis (IE) is a significant post-surgical challenge after cardiac surgery, characterized by severe bacterial invasion in the endocardium of the heart. To date, the most effective treatment strategy for IE involves the use of prophylactic antibiotics, which could effectively diminish bacterial infection in the injured valve or endocardium lining. ^27, 28^ In this study, we developed a smart bioelectric cardiac patch for sustained delivery of curcumin (Cur) under electrical stimulation to combat IE. We studied the pathogenesis of well-known gram-positive (MRSA, clinical isolate) and gram-negative (*E. coli*) bacteria as a model using the CSA, CSA4, and Cur-loaded CSA4 hydrogels for evaluating their antibacterial properties *in vitro*.

We observed a sustained release of Cur from CSA4 hydrogel under 250 mV (25 MHz) EF stimulation for up to 12 h. To verify the effect of Cur release and the effect of EFs dose on the antibacterial properties, we performed the plate dilution assay and the O.D.-based proliferation assay. The plates with PBS treatment were taken as controls.

As illustrated in **Fig. 3(a)**, the CSA4 + Cur group exhibited a negligible amount of *E. coli* and MRSA colonies compared to CSA and CSA4 w/o EFs stimulation. Besides, the EFs (250 mV, 25 MHz)-treated plates of *E. coli* and MRSA exhibited significantly fewer colonies in the control, CSA, and CSA4 groups. Interestingly, we observed no colony formation in both *E. coli* and MRSA following EF stimulation through CSA4 + Cur hydrogel, suggesting that EFs (250 mV, 25 MHz) itself had a positive role in reducing bacteria colony and Cur release from CSA4 hydrogel under EFs enhanced the colony reduction of both bacteria.

**Figure 3.**
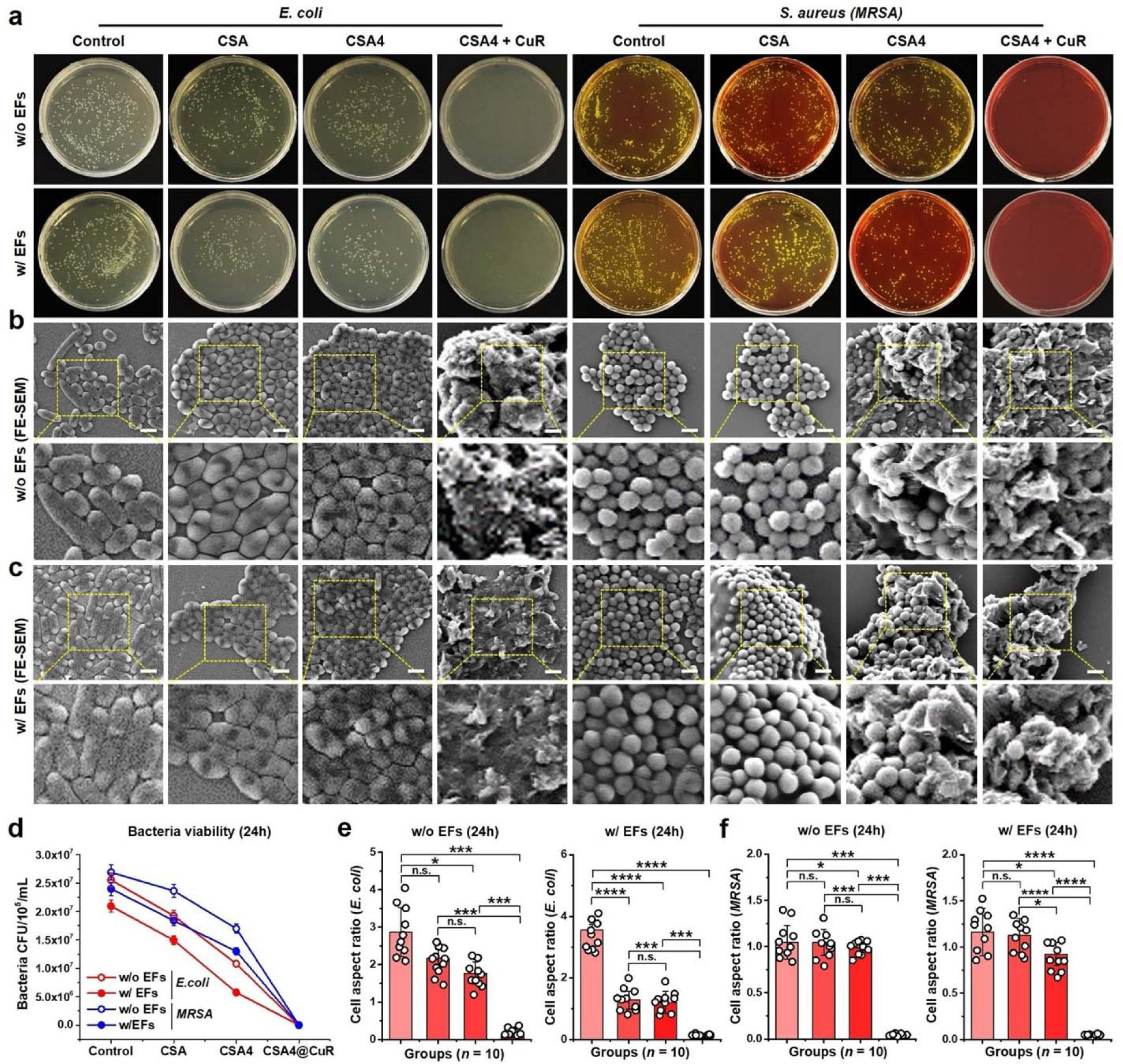
*In vitro* antibacterial performance of the nanoengineered adhesive hydrogels. **(a)** Digital photographs of the plate dilution test showing the bactericidal effect of the hydrogels w/ or w/o EFs stimulation against *E. coli* and *S. aureus* (MRSA) after 24 h of culture. The curcumin (CuR) loaded sample was tested to verify the synergistic effect of drug and EFs stimulation on bacteria. **(b, c)** Representative FE-SEM images of the bacteria showing morphological alteration and structural damage after 24 h of incubation. Scale bar: 2 µm. **(d)** The quantification of the colony forming unit (CFU) w/ or w/o EFs stimulation through the hydrogel platforms. **(e, f)** Calculation of the bacteria shape integrity (change in cell aspect ratio) w/ or w/o EFs stimulation as shown in the FE-SEM images. At least three (*N* = 3) independent images and ten (*n* = 10) bacteria/fields were used to assess the cellular integrity. Data represented as mean + s.d., statistical significance at \**p* < 0.05, \*\*\**p* < 0.001, and \*\*\*\**p* < 0.0001 (One-way ANOVA test).

To validate these findings, we next studied the ultrastructural morphology of bacteria using FE-SEM, and the results are shown in **Fig. 3(b, c)**. The control groups of *E. coli* exhibited smooth surface morphology w/o EFs. However, the CSA and CSA4 groups exhibited several damaged cells characterized by cell membrane rupture, demonstrating the antibacterial nature of chitosan (CS) and GO@ND. Notably, the CSA4 + Cur group showed significantly damaged *E. coli* cells, indicating the therapeutic efficacy of the drug-loaded hydrogel **(Fig. 3(b))**. Similarly, the control and CSA groups of MRSA displayed no noticeable damaged cells with mostly round morphology w/o EFs.

Besides, the CSA4 and CSA4+Cur groups exhibited mostly damaged cells characterized by ruptured cell membranes. More interestingly, the EF treatment significantly reduced the bacterial growth in both *E. coli* and MRSA, and Cur release synergistically improved the killing efficacy **(Fig. 3(c))**. Moreover, we also found a gradual reduction in bacterial colony-forming capability in synergistic treatment **(Fig. 3(d))**, demonstrating excellent antibacterial properties.

The quantification analysis of bacterial morphology was in good agreement with the plate assay results. As shown in **Fig. 3(e)**, the cell aspect ratio (long axis/short axis) disclosed a significant (***p < 0.001) reduction of *E. coli* cell body in CSA4 + Cur groups than other treated groups w/o or w/ EF stimulation. Similarly, the CSA4 + Cur group of MRSA also exhibited a greater reduction (***p < 0.001 and ****p < 0.0001) in cell aspect ratio **(Fig. 3(f))** than in the control and other groups, suggesting a potential change in morphology. Taken together, our results suggest that CS and GO@ND had a significant role in bacteria killing, which was significantly enhanced through EF stimulation and sustained Cur release from CSA4 hydrogel. Compared to previous reports, the developed electroactive hydrogel patch with smart drug delivery could be beneficial for treating post-surgical endocarditis in clinical settings.

### Effect of electromechanical stimulation on macrophage polarization

**Fig. 4(a)** schematically demonstrates the macrophage polarization study using 3D-printed anisotropic scaffolds. Prior to the polarization study, we tested the biocompatibility of RAW 264.7 cells using the WST-8 assay. As shown in **Fig. 4(b)**, the CSA4 hydrogel exhibited good biocompatibility with RAW 264.7 cells. It displayed significantly (*p < 0.05) higher viability than the control and CSA hydrogels w/o EF stimulation after 24 h of incubation, indicating its biocompatibility. However, no significant increase in viability was found in the CSA group when compared to the control. Strikingly, under EF stimulation, the CSA4 group showed more significant (*p < 0.05) viability than other groups **(Fig. 4(c))**, suggesting that electro-mechanotherapy is safe for the RAW 264.7 cells. To verify these findings, we further evaluated the viability using a live/dead assay, and the results are shown in **Fig. 4(d)**. Notably, neither CSA nor the CSA4 scaffolds were found cytotoxic in the presence or absence of EFs after 24 h of culture. The percentage of live cells was found to be significantly (***p < 0.001) higher in both groups **(Fig. 4(e))**, suggesting their superior biocompatibility. Thus, consistent with our previous report ^6^, we suggest that 250 mV/20 min EF stimulation is biologically safe along with the conductive CSA hydrogel.

**Figure 4.**
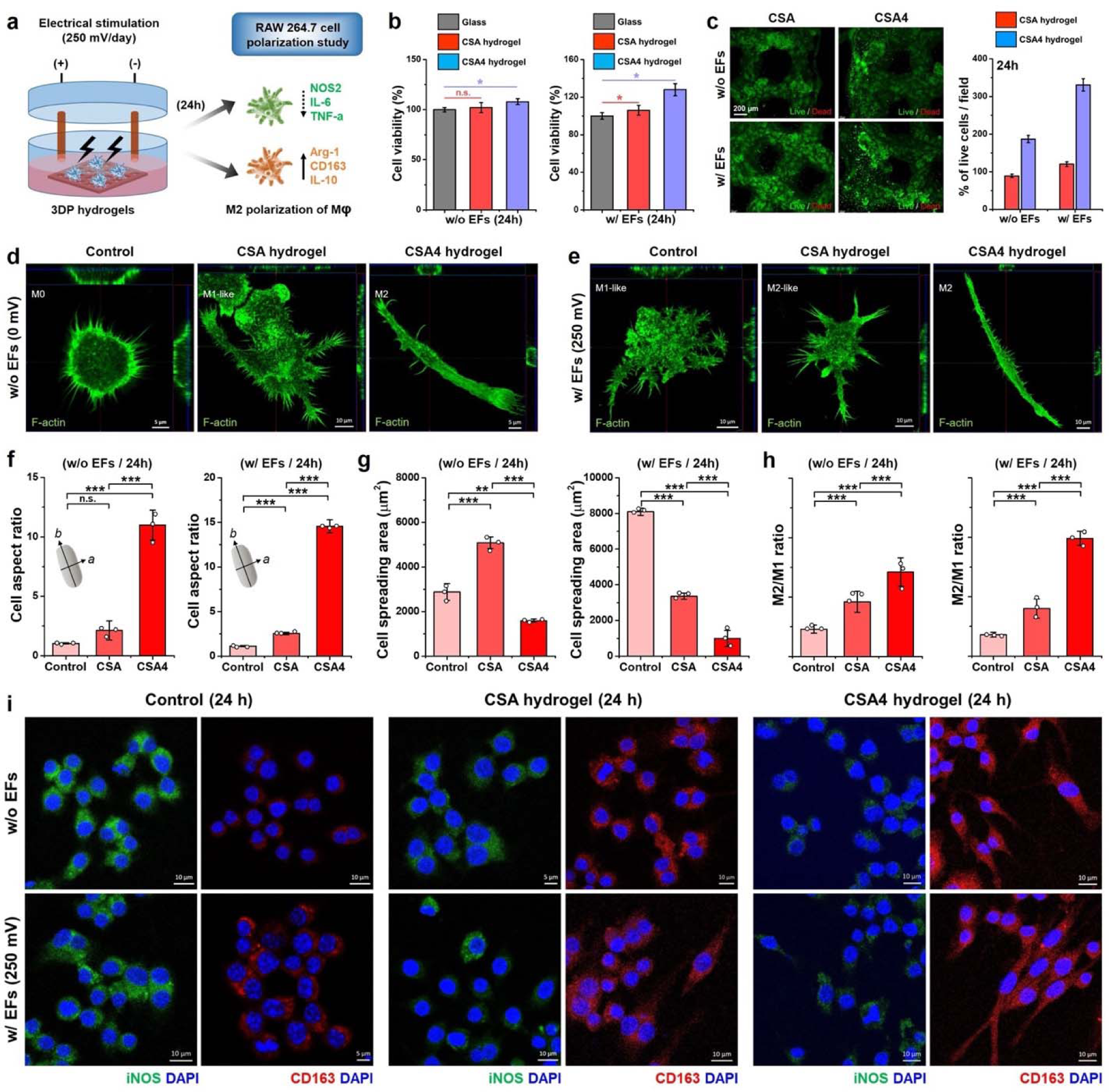
Macrophage polarization potential of the developed scaffolds after 24 h w/ or w/o EFs in the presence of 3D-printed anisotropic scaffolds. **(a)** Schematic illustration of the macrophage polarization study. **(b)** WST-8 assay of RAW 264.7 cells with fabricated scaffold w/ or w/o EFs at indicated time point. **(c)** Representative live/dead assay of RAW 264.7 cells with CSA and CSA4 scaffold in the presence or absence of EFs with quantification data. Scale bar: 200 µm. **(d, e)** CLSM images with orthogonal cut images showing the volumetric F-actin expansion in various groups w/ or w/o EFs stimulation after 24 h. Scale bar: 5 and 10 µm **(f, g)** Calculation of the cell aspect ratio and spreading area of RAW 264.7 cells in various groups. **(h)** Calculation of macrophage polarization in various groups. **(i)** CLSM images of the RAW 264.7 cells showing the expression of iNOS (M1 phenotype) and CD163 (M2 phenotype) markers after 24 h of culture. Scale bar: 5, 10, and 20 µm. Data represented as mean ± s.d. of triplicate (*n* = 3) experiments, statistical significance considered at \**p* < 0.05, \*\**p* < 0.01, and \*\*\**p* < 0.001 with One-way ANOVA test.

To investigate the effect of electromechanical stimulation on RAW 264.7 cell polarization, we examined the macrophage morphology using confocal laser scanning microscopy (CLSM) and immunocytochemistry (ICC) after 24 h of incubation. As shown in **Fig. 4(f)**, the control RAW 264.7 cells exhibited round morphology (dendritic, M0) with non-branched filopodia w/o EF stimulation. The cells grown on CSA hydrogel exhibited mostly oval to ‘poached egg’-like morphology with branched filopodia, suggesting typical M1-type polarization. Interestingly, the cells grown on CSA4 hydrogel showed a flat and elongated (fibroblastic) morphology with reduced filopodia or lateral branching, resembling the M2 polarization. The elongation of macrophages was probably owing to the lamellar and aligned structure with the porous geometry of the CSA4 scaffold, which facilitated the mechanical activation during polarization. ^8, 29^ Conversely, we observed flat and fibroblastic cell morphology in the CSA and CSA4 groups under EF stimulation (250 mV, 50 kHz), suggesting the M2-like and M2 phenotypes of macrophages **(Fig. 4(g))**. Besides, the control group exhibited a typical M1 phenotype even after EF stimulation for 24 h. To verify these findings, we then studied the cell morphological parameters. As illustrated in **Fig. 4(h)**, a significantly (***p < 0.001) higher cell aspect ratio was noted in the CSA4 group than in other groups, w/ or w/o EF stimulation. Meanwhile, the cell spreading area was significantly reduced in CSA4 groups than in control and CSA groups, w/ or w/o EFs, indicating the change in cell shape and morphology **(Fig. 4(i))**. The quantification of macrophage polarization ratio (M2/M1) in various treatment groups is shown in **Fig. 4(j)**.

To validate the macrophage polarization, we further examined the intracellular localization of pro-inflammatory (iNOS) and anti-inflammatory (CD163) markers through ICC after 24 h of incubation. As demonstrated in **Fig. 4(k)**, the control groups displayed predominantly higher iNOS (∼95%) and negligible CD163 (∼1%) expression w/o EFs treatment. The CSA group showed moderate iNOS (∼57%) and CD163 (∼60%) expression. Notably, the CSA4 group exhibited low iNOS (∼12%) and higher CD163 (∼88%) expression after 24 h of incubation, w/o EFs. More interestingly, after EF stimulation, the CSA4 groups exhibited a higher amount of CD163^+^ (∼92%) and iNOS^-^ (∼3%) cells than the control and CSA groups, suggesting that synergistic electro-mechanotherapy had a significant role in M2 polarization. The anti-inflammatory effect of CSA4 partially comes from the GO complex, ^30^ which has been shown to induce an immune response in macrophages. It has been reported that alternating EF and direct EF-responding biomaterials^6^ had better M2 polarization efficiency than piezoelectric materials^5^, where the M2/M1 polarization was facilitated by enhanced expression of pro-inflammatory factors by elevating the potassium voltage-gated ion channels (*KCNC1* and *KCNH2*), respectively. Gu et al.^31^ reported that sinusoidal EF induces M2 macrophage polarization via upregulating the expression of CD206, IL-10, and TLR4 transcription factors. Consistent with previous reports, we concluded that the sinusoidal EFs with aligned topography of the conductive CSA4 scaffold facilitated the M2 macrophage polarization, which would be beneficial for healing the damaged cardiac tissue, especially during MI or post-valvular infections.

### Scaffold’s electromechanical cues regulate hCM viability and orientation

To investigate the synergistic role of electrical stimulation and scaffold stiffness on hCM’s biomechanical fate, we first performed microindentation-based measurements. **Fig. 5(a)** represents a comparison of the concentration-based stiffness of some commonly used ECM-mimicking hydrogels vs. fabricated hydrogels used in this study. Interestingly, the Differential modulus (*K*’) was significantly (***p < 0.001) increased as the concentration of GO@ND increased in the CSA matrix. Among all the fabricated hydrogels, the CSA4 displayed significantly (***p < 0.001) higher stiffness than the control and other groups **(Fig. 5(b))**. Since cardiac impulses and contractions require a stiffness of <10 kPa (typically <500 Pa), we fabricated all hydrogels below that range. The stiffness for CSA, CSA1, CSA2, and CSA4 scaffolds was calculated to be ∼1.6 kPa, 3.3 kPa, 4.1 kPa, and 6.2 kPa, respectively. FE-SEM images of the hydrogel scaffolds with varying stiffness and topography used for the hCMs orientation and differentiation are shown in **Fig. 5(c)**.

**Figure 5.**
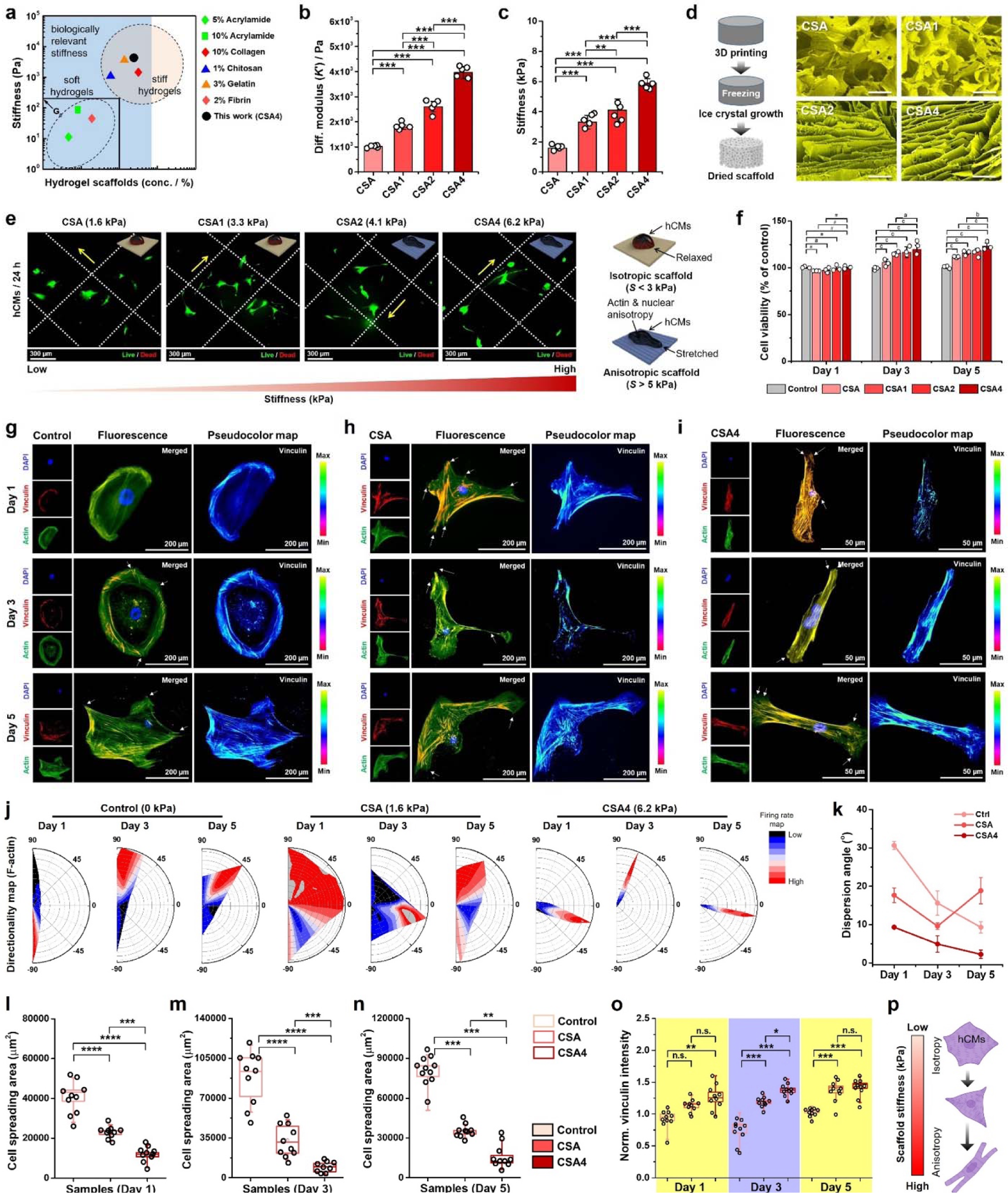
Effect of scaffold topography (stiffness) on human cardiomyocytes (hCMs) proliferation and adhesion. **(a)** A comparative study of the stiffness of various ECM-mimicking biopolymers was reported in the literature. **(b, c)** Microindentation results (*n* = 5) showing the differential modulus (*K*′, Pa) and stiffness (*S*, kPa) of the fabricated nanoengineered hydrogels. **(d)** Schematic illustration of the 3D printed hydrogel scaffold preparation with corresponding SEM morphologies showing the anisotropic pore distribution. Scale bar: 5 µm. **(e)** Representative fluorescence microscopy images of the hCMs showing the live and dead cells growing onto the 3D-printed scaffolds with varying stiffness (1.6-6.2 kPa) after 24 h of culture. Scale bar: 300 µm. **(f)** WST-8 cell viability results of hCMs growing onto the 3D-printed scaffolds at indicated time points. Data reported here are mean ± s.d. of triplicated (*n* = 3) experiments, statistical significance at \**p* < 0.05 [*a*], \*\**p* < 0.01 [*b*], and \*\*\**p* < 0.001 [*c*], analyzed through Student *t*-test. **(g-i)** Immunostaining images of hCMs showing the expression of vinculin (red) with cytoskeleton (F-actin, green) in various groups. Scale bar: 50 and 200 µm. **(j)** Order parameters with corresponding angular distributions (directionality) and firing rates (*n* = 10) of the F-actin in various groups showing the cellular anisotropy. **(k)** Dispersion parameters of the F-actin in degrees at indicated time points. **(l-n)** Representative calculation of cell spreading area (µm^2^) of hCMs cultured onto the fabricated scaffolds with various stiffness at indicated time points. At least ten (*n* = 10) independent images were randomly selected for ImageJ measurements. Data reported here are mean ± s.d., statistical significance at \*\**p* < 0.01, \*\*\**p* < 0.001, and \*\*\*\**p* < 0.0001 (Tukey’s HSD *post-hoc* analysis). **(o)** Quantification of vinculin intensity of hCMs under various treatment conditions. Data reported here are mean ± s.d., statistical significance at \**p* < 0.05, \*\**p* < 0.01, and \*\*\**p* < 0.001 (Tukey’s HSD *post-hoc* analysis). **(p)** Schematic diagram showing the effect of scaffold stiffness and anisotropy on hCMs orientation.

The *in vitro* biocompatibility of the fabricated anisotropic scaffolds with varying stiffness (1.6–6.2 kPa) on hCMs was studied using live/dead and WST-8 assays after 24 h of culture. As depicted in **Fig. 5(e)**, the hCMs were found to be highly biocompatible with the 3D scaffolds. The hCMs were found to be well-adhered to the scaffold surface, regardless of their stiffness. However, the hCMs grown on CSA2 and CSA4 scaffolds showed elongated morphology with many filopodia, which was probably due to the anisotropic geometry of the scaffold’s pore. Based on this observation, we hypothesized that soft and isotropic scaffolds (less or no GO@ND content) with stiffness ∼3 kPa showed regular cell morphology (relaxed state), while anisotropic scaffolds (high GO@ND content) with stiffness > 3 kPa displayed elongated (stretched) hCMs, respectively. To verify this finding, we performed the WST-8 assay, and the result is shown in **Fig. 5(f)**. On day 1, all the scaffolds showed no significant change in hCMs viability. However, on day 3, the CSA1, CSA2, and CSA4 groups exhibited a significant enhancement (***p < 0.001) in viability (>110%) compared to the control group. Furthermore, the CSA4 groups displayed a more significant (**p < 0.01) increase in viability (∼122%) than the control after 5 days of incubation, suggesting that CSA scaffolds with varying stiffness (1.6–6.2 kPa) positively regulate the viability of hCMs.

Based on the outstanding viability result, we further studied the mechanisms of hCM adhesion and orientation up to 5 days of culture, and the results are shown in **Fig. 5(g-i)**. It is important to note that we compared the cell adhesion dynamics of CSA4 with control (0 kPa) since the highest cell viability was found in the presence of CSA4 (GO@ND content: 4 wt.%, high stiffness). Besides, we have also taken the CSA scaffold (devoid of GO@ND, low stiffness) as a positive control for this experiment. It has been reported that vinculin and metavinculin expressions are critical during force transduction at the focal adhesion sites (FAs) in cells. ^32, 33^ Understanding the fact that FA kinases (FAKs) are closely associated with cytoskeletal proteins during cell migration ^34^, we studied the expression profile of Vinculin and F-actin orientation in response to CSA scaffold anisotropy and stiffness. **Fig. 5(g)** represents the CLSM image with the corresponding pseudocolor map for vinculin of hCM growing on TCPS at indicated time points. In the control group (TCPS), we observed a stable expression of vinculin, mostly distributed to the cell periphery with F-actin on days 1 and 3. On day 5, the Vinculin expression was spotted on the protruding filopodia at a higher rate than distributed locally, indicating that cell migration affected the differentiation expression of FAK.

Interestingly, this phenomenon was more prominent when the cells were grown on the CSA scaffold **(Fig. 5(h))**. In the case of the CSA scaffold (stiffness: 1.6 kPa), higher vinculin expression was noted at or near the FA sites, while scanty expression was observed in the cytoplasmic region, as evidenced through pseudocolor imaging.

Remarkably, the anisotropic CSA4 scaffold (stiffness: 6.2 kPa) displayed an inverse pattern of Vinculin expression, as shown in **Fig. 5(i)**. As the hCM was found to elongate from day 1 to day 5, a gradual decrease in filopodia formation and subsequent vinculin expression was spotted. However, a lesser amount of vinculin was spotted along with peripherally aligned F-actin fibers, suggesting that the scaffold anisotropy and stiffness had a positive role in FAK expression. No significant change in vinculin expression was noted in the CSA and CSA4 groups. The greater expression of vinculin in the CSA scaffold was probably due to the random porosity and less mechanical forces exerted by the cells. In contrast, the CSA4 scaffold with anisotropic geometry transduces greater biomechanical forces, resulting in a force-driven change in morphology, which was consistent with previous reports. ^32^ The normalized vinculin intensities of various groups are given in **Fig. 5(o)**.

The F-actin orientation and morphometric results also agree with the FAK expression. As shown in **Fig. 5(j)**, the directionality map showed a random distribution of F-actin fibers in the control group for up to 5 days. However, in the CSA group, most F-actins were found randomly up to 3 days, and on day 5, most of the cells exhibited aligned F-actins. Besides, owing to the anisotropic and higher forces exerted by the CSA4 scaffold, all the cells were found to display anisotropic distribution of F-actin for up to 5 days. This was also confirmed by measuring the dispersion angle, as shown in **Fig. 5(k)**. We also observed a change in the cell spreading area, supporting the previous findings. On day 1, the spreading area was significantly (***p < 0.001) decreased in CSA4 group cells compared to other groups **(Fig. 5(l))**. Moreover, this changing pattern was predominantly spotted on day 3 **(Fig. 5(m))** and day 5 **(Fig. 5(n))** for all the groups, suggesting that scaffold anisotropy and stiffness-assisted alignment significantly change the cell spreading area while maintaining a low amount of vinculin expression. The nuclear spreading area was weakly correlated (y = 0.014 + 277.9, R^2^ = 0.1679) with the cell spreading area for all the groups without EFs, suggesting that scaffold anisotropy and stiffness had no direct correlation with nuclear changes. A schematic illustration of scaffold biomechanical force-driven cellular anisotropy is demonstrated in **Fig. 5(p)**.

### Cellular anisotropy confers hCMs differentiation under electromechanical stimulation

Based on the findings mentioned above, we next applied the EFs (250 mV, 20 min/day) to the hCMs growing on scaffolds. The cardiomyogenic differentiation was observed after 5 days of incubation with various scaffolds (stiffness: 0, 1.6, and 6.2 kPa), w/ or w/o EF stimulation. The control group exhibited stable expression of a-actinin (a-ACT) and cardiac troponin T (cTnT) in w/o or w/ EFs treatment **(Fig. 6(a))**. The CSA (1.6 kPa) and CSA4 (6.2 kPa) groups showed no noticeable change in a-ACT and cTnT expression. Strikingly, the expression profile for smooth muscle actinin (SMA) and cardiac troponin 1 (cTn1) was found to be significantly higher in the CSA and CSA4 groups **(Fig. 6(b))** than in control, suggesting that cardiac cell differentiation was regulated through scaffold stiffness and EF stimulation. More interestingly, we found that hCMs growing on CSA and CSA4 with EFs displayed more prominent nuclear and cytoplasmic expression of N-cadherin (NCAD) than EFs-untreated groups **(Fig. 6(c))**.

**Figure 6.**
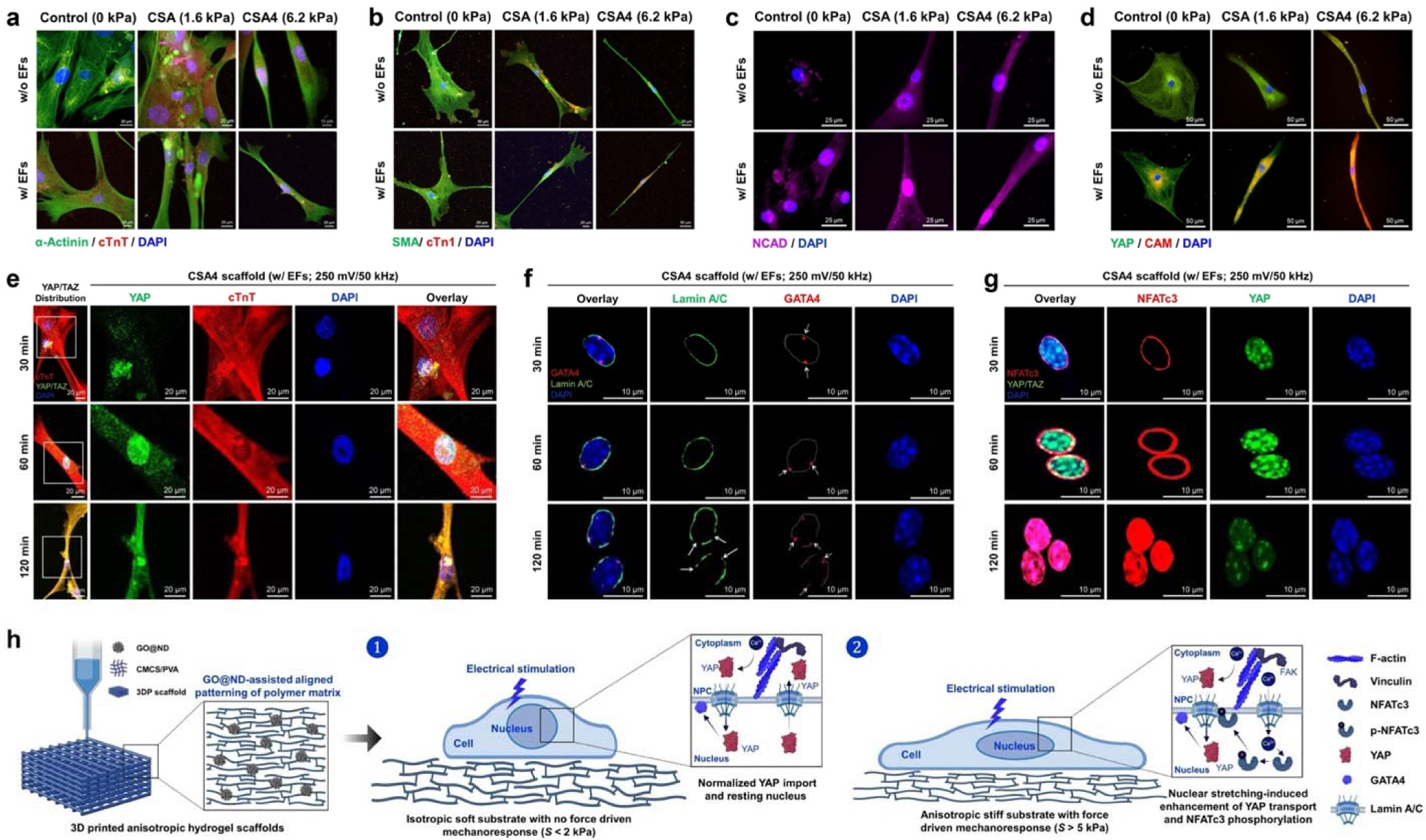
Electromechanical force-driven nuclear trafficking of YAP and activation of GATA4/NFATc3 is crucial during hCMs differentiation. **(a-d)** Representative immunofluorescence images of hCMs showing the expression profile for α-actinin/cTnT (green/ red), SMA/cTn1 (green/red), NCAD (magenta), and YAP/CaM (green/red) with nucleus (DAPI, blue) growing on scaffolds with varying stiffness (0, 1.6, and 6.2 kPa). Scale bar: 20 and 25 µm. **(e)** Time-dependent (30-120 min) activation of YAP (green) against cTnT (red) and its intracellular/extracellular transport is regulated by EFs stimulation and CSA4 scaffold stiffness. Scale bar: 20 µm. **(f)** Time-dependent Lamin-A/C (green) degradation confers transport of GATA4 (red) towards the nuclear membrane. Scale bar: 10 µm. **(g)** Representative immunofluorescence imaging of hCMs showing the time-dependent nuclear expression of NFATc3 (red) and YAP (green) protein, post-EF stimulation on CSA4 scaffold. Scale bar: 10 µm. **(h)** A hypothetical mechanism of scaffold anisotropy and electromechanical force-driven mechanotransduction in hCMs.

Also, the cytoplasmic localization of the yes-associated protein (YAP) and calmodulin (CaM) was found to be exceptionally higher in the CSA4 (6.2 kPa) scaffold group w/ EF stimulation **(Fig. 6(d))**, suggesting that the electro-mechanical force-driven hCM differentiation is dependent on actin anisotropy, vinculin activation, and YAP localization.

### hCMs respond to electromechanical stress via phosphorylating NFATc3 through YAP/TAZ and Lamin-A/C mediated nuclear trafficking

It has been reported that surface topography has a significant impact on cellular mechano-transduction in cardiac cells, which is principally governed by YAP/TAZ activation and its nuclear transport. ^35, 36^ Thus, inhibition of YAP/TAZ signaling attenuates its nuclear transport and hinders the activation of the NFATc3 nuclear factor and subsequent reduction of hCM development. ^37^ Herein, we investigated the electromechanical cues associated with hCMs differentiation through a 3D-printed conductive CSA4 scaffold (stiffness: 6.2 kPa) under sinusoidal EFs (250 mV, 50 kHz) stimulation in a time-dependent manner. We examined the activity of mechanosensitive cytoplasmic and nuclear factors within a time frame of 30-120 min post-stimulation. **Fig. 6(e)** illustrates the time-dependent localization of YAP in hCMs within a time frame of 30–120 min. It was interesting to note that after 30 min post-stimulation, the nuclear YAP localization was spotted with a scanty expression in the nucleus. However, the nuclear and cytoplasmic fluorescence of YAP remarkably increased after 60 min. Furthermore, we also observed a gradual increase of nuclear YAP at 120 min post-stimulation, indicating that nuclear import of YAP is not only governed by EF stimulation but also due to substrate stiffness. ^38, 39^

To verify this finding, we then investigated the nuclear transport machinery and chromatin remodeling involved with CSA4+EF stimulation by ICC analysis, and the results are shown in **Fig. 6(f)**. The CLSM images confirmed that 30 min post-stimulation induced the loss of Lamin-A/C and was predominant up to 120 min of observation. Also, the loss of Lamin-A/C integrity activated the early accumulation of GATA4. After 60 min post-stimulation, the Lamin-A/C boundary was clearly observed, following a distant localization of the GATA4 transcription factor. Interestingly, at around 120 min post-stimulation, the loss of Lamin-A/C integrity was clearly visible in the close vicinity of GATA4, confirming the activity of the nuclear pore complex (NPC). ^38^ This phenomenon can be correlated with the CSA4 scaffold’s anisotropy and stiffness, which facilitated the average nuclear stretching and subsequent YAP transport. ^40^ Besides, we also found that the gradual import of YAP **(Fig. 6(g))** enhanced the phosphorylation of NFATc3 around 60 min post-stimulation on the CSA4 scaffold, confirming that stiffer matrix and EF stimulation synergistically enhanced the nuclear YAP localization within 120 min and thereby regulating the expression of cTnT during cardiomyogenesis.

## CONCLUSION

In summary, we presented a multifunctional 3D printable hydrogel patch with controllable stiffness for cardiac tissue engineering. The as-fabricated nanoengineered hydrogel displayed robust mechanical and adhesive properties with good sensing abilities and was suitable for cardiac bioelectronics fabrication. Besides, the fabricated hydrogels showed excellent biocompatibility with RAW 264.7 and showed anti-inflammatory activation under sinusoidal EF (250 mV/20 min/day) stimulation. In hCM, the CSA4 scaffold with a stiffness of 6.2 kPa displayed early activation of YAP/TAZ under electromechanical stress, which later induced phosphorylation of NFATc3 and subsequent trafficking through Lamin A/C during cardiomyogenesis. We hope that this work will enlighten future research on the electro-mechanobiology of cardiac tissues for better management of arrhythmic and infracted heart disease.

## METHODS

### Fabrication of nanoengineered hydrogel patch

The carboxymethyl chitosan (CMCS) was synthesized according to our previous report ^23^ and characterized using ^1^H-NMR and FT-IR spectroscopy. The GO@ND nanocomposite was synthesized as reported elsewhere. For hydrogel fabrication, the desired amount of PVA (Daejung) and CMCS was dissolved in D.I. water (Millipore) under vigorous stirring to achieve a final concentration of 10% (w/v) and 5% (w/v). After that, the GO@ND was mixed with respect to the weight of CMCS/PVA and stirred overnight to obtain a homogenous mixture. The concentration of GO@ND used in this study was 1-4 wt.%, respectively. Next, 2 wt.% borax (Sigma-Aldrich) was added to the hydrogel mixture to initiate borate crosslinking. For 3D printing, the hydrogels were prepared without borax and loaded onto the printing cartridges.

### 3D printing and fabrication of anisotropic scaffolds

The 3D printing was carried out using a Cellink BioX (Cellink, Sweden) 3D bioprinter. The printing parameters and processes are listed in our previous study. ^6^ After 3D printing, the hydrogels were slightly crosslinked with a 0.5% glutaraldehyde (Sigma-Aldrich) solution for 10 min to maintain their structural integrity. After chemical crosslinking, the hydrogels were freeze-dried to obtain anisotropic scaffolds with varying stiffness.

### General characterizations

The morphology of the GO@ND nanocomposite was studied using FE-SEM (Jeol), HR-TEM (Jeol), and AFM (Nanoscope V). A rotational rheometer (Anton Paar) analyzed the viscoelastic nature and shear-thinning properties. An FE-SEM (Jeol) observed the scaffold geometry and ultrastructure. The scaffold’s stiffness was measured using the microindentation (UNHT^3^ Bio, Anton Paar) technique. The electrical conductivity and electrochemical properties were evaluated using a source meter (Keithley) connected to a 4-probe (PASCO) station. The sensing performance of the hydrogel film was probed through a high-performance digital source meter (Keithley). The mechanical and adhesive behavior of the hydrogel films were monitored using a universal testing machine (UTM, A&D Digital). The *in vitro* bioadhesive test was performed using porcine skin and murine heart samples.

### Drug loading and antibacterial studies

Curcumin (Sigma-Aldrich) was taken as a model drug for studying the release profile under EF stimulation and subsequent antibacterial properties. The pathogenic bacteria Methicillin-resistant *Staphylococcus aureus* (MRSA) and *Escherichia coli* (*E. coli*) were selected as model gram-positive and gram-negative organisms for antibacterial studies. The antibacterial efficacy was assessed using optical density (OD) measurements, agar plating techniques, and SEM analysis. Initially, MRSA was cultured in tryptic soy broth and *E. coli* in nutrient broth at 37 ℃ for an overnight period, resulting in the production of a new batch of bacteria.

Subsequently, the optical density (OD) of the newly prepared culture was measured at a wavelength of 600 nm and then adjusted to a value of 0.1 (100μl) before commencing the experiment. An equal amount of samples was cultured with bacteria w/ and w/o electrical stimulation (250 mV, 25 MHz) for 12 h. Optical density (OD) was measured, and agar plating was conducted using a dilution factor of 10^-5^ to assess the impact of CSA hydrogels and ES. The bacteria were further treated with 4% paraformaldehyde for fixation and then dehydrated using a series of ethanol concentrations (10%, 30%, 50%, 70%, 80%, 90%, and 100%). Scanning electron microscopy (SEM) was used to capture pictures for morphological analysis.

### Macrophage viability and polarization study

Macrophage viability was assessed using WST-8 (Cellrix) and Live/Dead (Sigma-Aldrich) staining assays after 24 h of incubation on scaffolds w/ or w/o EF stimulation (250 mV, 50 kHz, 20 min/day). The EFs dose adopted here was reported in our previous study. ^20^ The macrophage polarization study was conducted using F-actin staining and immunofluorescence analysis. For morphological staining, RAW 264.7 (∼1.5 x 10^4^) cells were fixed with 3.7% PFA (Sigma-Aldrich) for 15 min, followed by permeabilization with 0.1% TX-100 (Sigma-Aldrich) for 10 min and blocking with 1% BSA (Sigma-Aldrich) for 1 h. Next, cells were stained with an F-actin probe (Thermo-Fischer), and the nucleus was counterstained with DAPI (Sigma-Aldrich). For immunofluorescence, RAW 264.7 (∼2.5 x 10^4^) cells were fixed, permeabilized, and blocked with standardized protocols. After that, the cells were incubated with the appropriate primary and secondary antibodies (Santa Cruz Biotechnology), followed by DAPI staining. Cells were visualized using an inverted confocal microscope (LSM, Zeiss) with appropriate filters.

### Human cardiomyocyte culture and electro-mechanostimulation

For electro-mechanostimulation in the presence of 3D conductive scaffolds, human ventricular cardiomyocytes (hCMs, ATCC) were cultured in myocyte growth media. The viability of hCMs was examined using a Live/Dead staining kit and the WST-8 assay after the desired time intervals. For studying cellular anisotropy and F-actin orientation, the hCMs (∼2.5 x 10^4^) were cultured on CSA (stiffness: 1.6 kPa) and CSA (stiffness: 6.2 kPa) scaffolds for 5 days. After that, the cells were fixed, permeabilized, and blocked with strand protocols. Next, the cells were incubated with a primary antibody against Vinculin (Santa Cruz Biotechnology), followed by incubation with a fluorescent secondary antibody. After that, the cytoskeleton was stained with F-actin (Thermo-Fischer), followed by nucleus staining with DAPI (Sigma-Aldrich). The hCM cell adhesion and orientation were observed using an inverted fluorescence microscope (DMi8, Leica).

### Immunocytochemistry

The effect of electro-mechanostimulation on hCM differentiation and the underlying mechanosensing was investigated using an immunofluorescence study after 5 days of culture. The nuclear mechanosensing study was performed within a time frame of 30-120 min post-stimulation on the CSA4 scaffold. For this, hCMs (∼2.5 x 10^4^) were fixed with 3.7% PFA (Sigma-Aldrich) for 15 min, followed by permeabilization with 0.5% TX-100 (Sigma-Aldrich) for 10 min. Next, the cells were blocked with 1% BSA for 1 h. After that, the cells were incubated with appropriate primary and secondary antibodies (Abcam and Santa Cruz Biotechnology) overnight at 4 °C. Next, the cells were washed with PBS (Welgene) and stained with DAPI (Sigma-Aldrich). Finally, cells were mounted (Invitrogen) and visualized using an inverted confocal microscope (Zeiss) with appropriate lasers.

### Image analysis

For confocal and fluorescence microscopy, the images were acquired using ZEN lite (Zeiss) and LAS-X (Leica) software. The images were analyzed using ImageJ (v1.5, NIH) software. The cell orientation and directionality maps were analyzed using ImageJ Fiji (v1.5.4, NIH) with the ‘Directionality’ plugin (Java v1.8.0) and plotted using Prism (GraphPad) software.

### Statistical analysis

The statistical analysis was carried out using Origin Pro (v9.1). The significant difference between the control and treatment groups was calculated using a One-way Analysis of Variance (ANOVA), followed by a student *t*-test and Tukey’s HSD *post-hoc* analysis. The statistical significance was considered at *p < 0.05, **p < 0.01, ***p <0.001, and ****p < 0.0001.

## AUTHORS CONTRIBUTION

S.D.D. and T.V.P. contributed equally to the manuscript. S.D.D. wrote the manuscript. T.V.P. and K.T.L. revised the manuscript. All the authors read and approved the final draft of the manuscript.

## ACKNOWLEDGMENTS

We express our sincere gratitude to the lab members for their kind suggestions during the manuscript preparation. We thank Ms. Rumi Acharya and Mr. Hojin Kim for their kind contribution to hydrogel preparation. This study was supported by the ‘Basic Science Research Program’ through the ‘National Research Foundation of Korea’ funded by the ‘MoE’ (NRF-2018R1A16A1A03025582; NRF2022R1I1A3063302). This work was supported by the Innovative Human Resource Development for Local Intellectualization program through the Institute of Information & Communications Technology Planning & Evaluation (IITP) grant funded by the Korean Government (MSIT) (IITP-2024-RS-2023-00260267).

## DATA AVAILABILITY

The data is made available upon request.

## RESOURCE TABLE

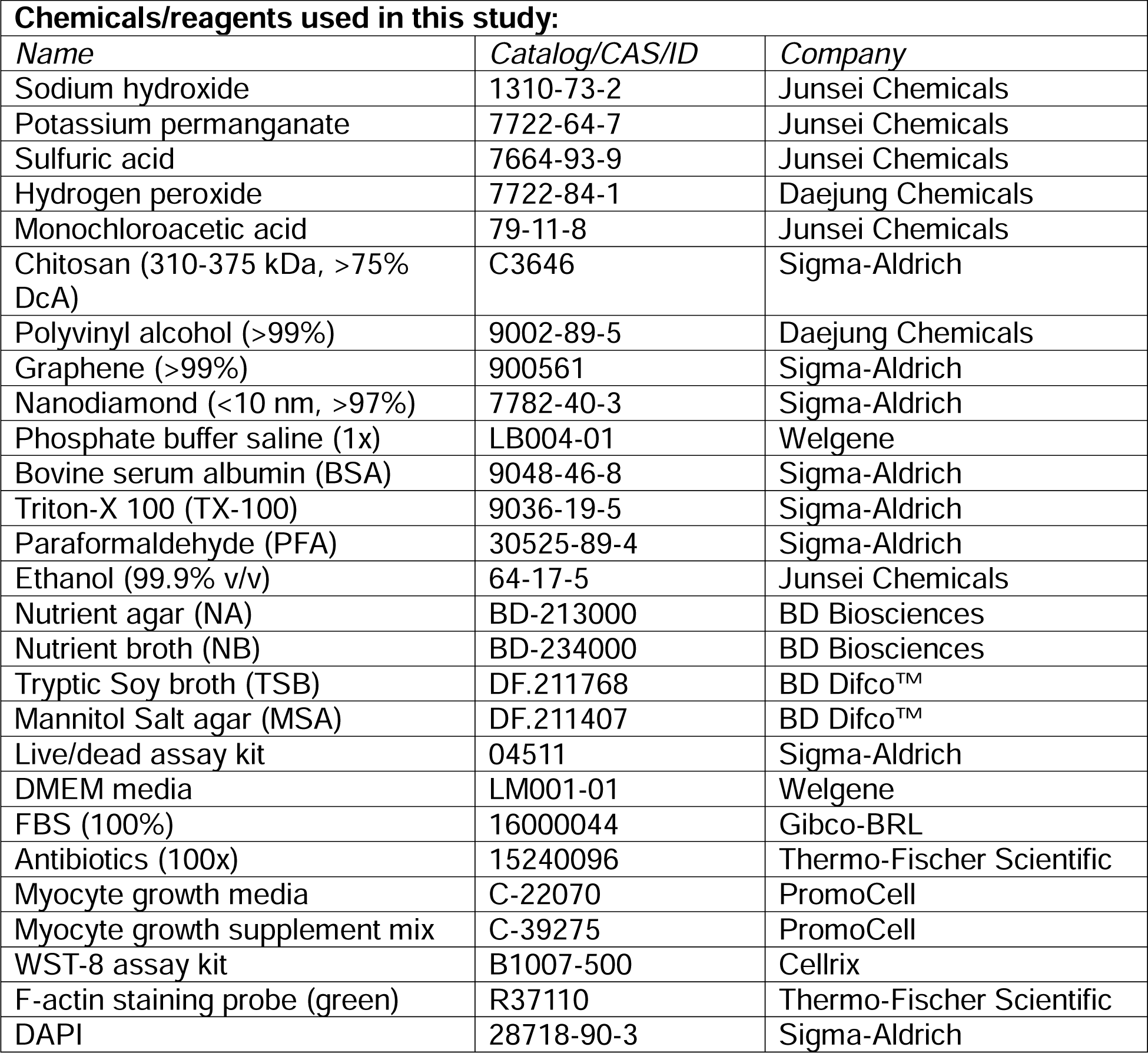
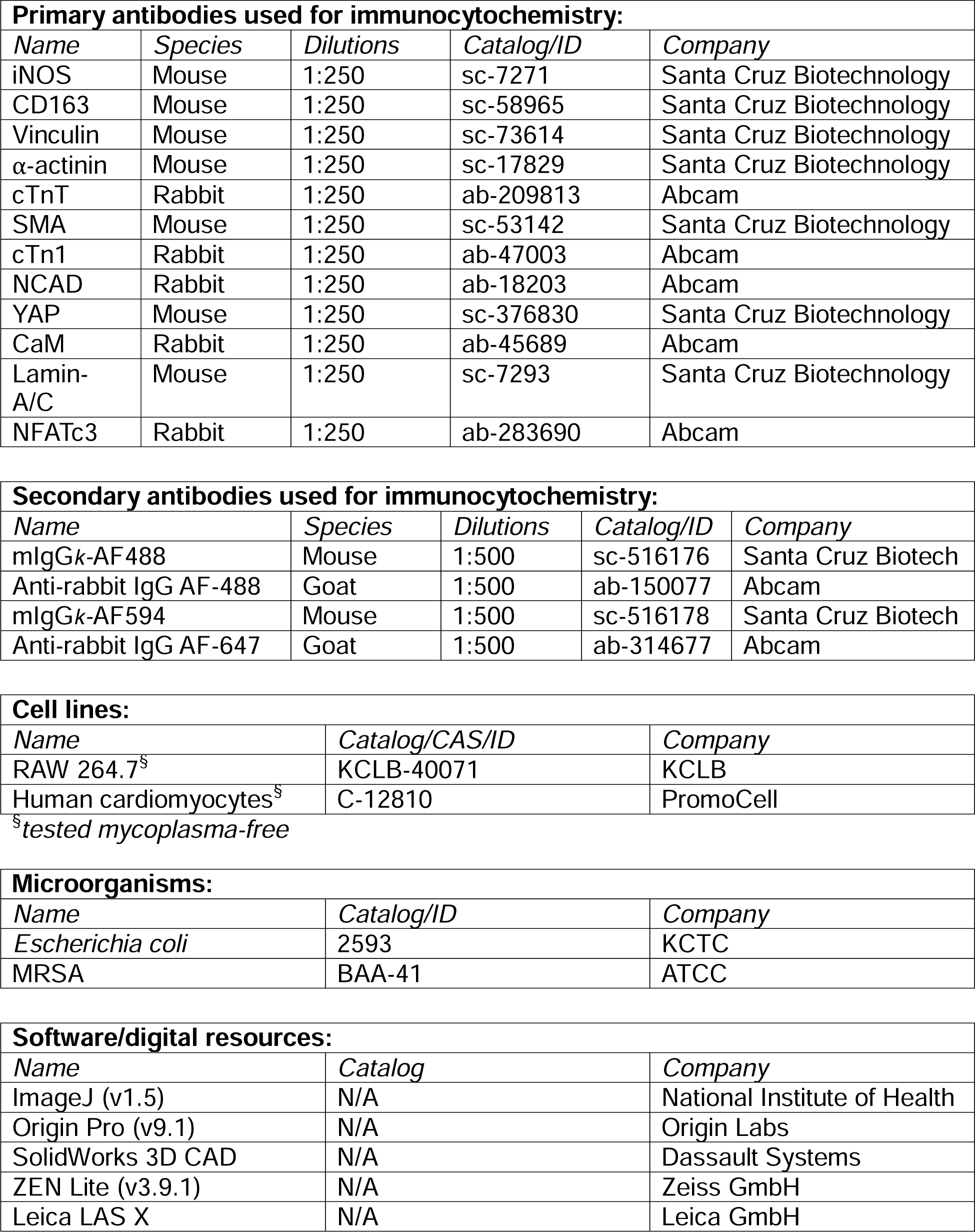

---

**Scheme 1.**
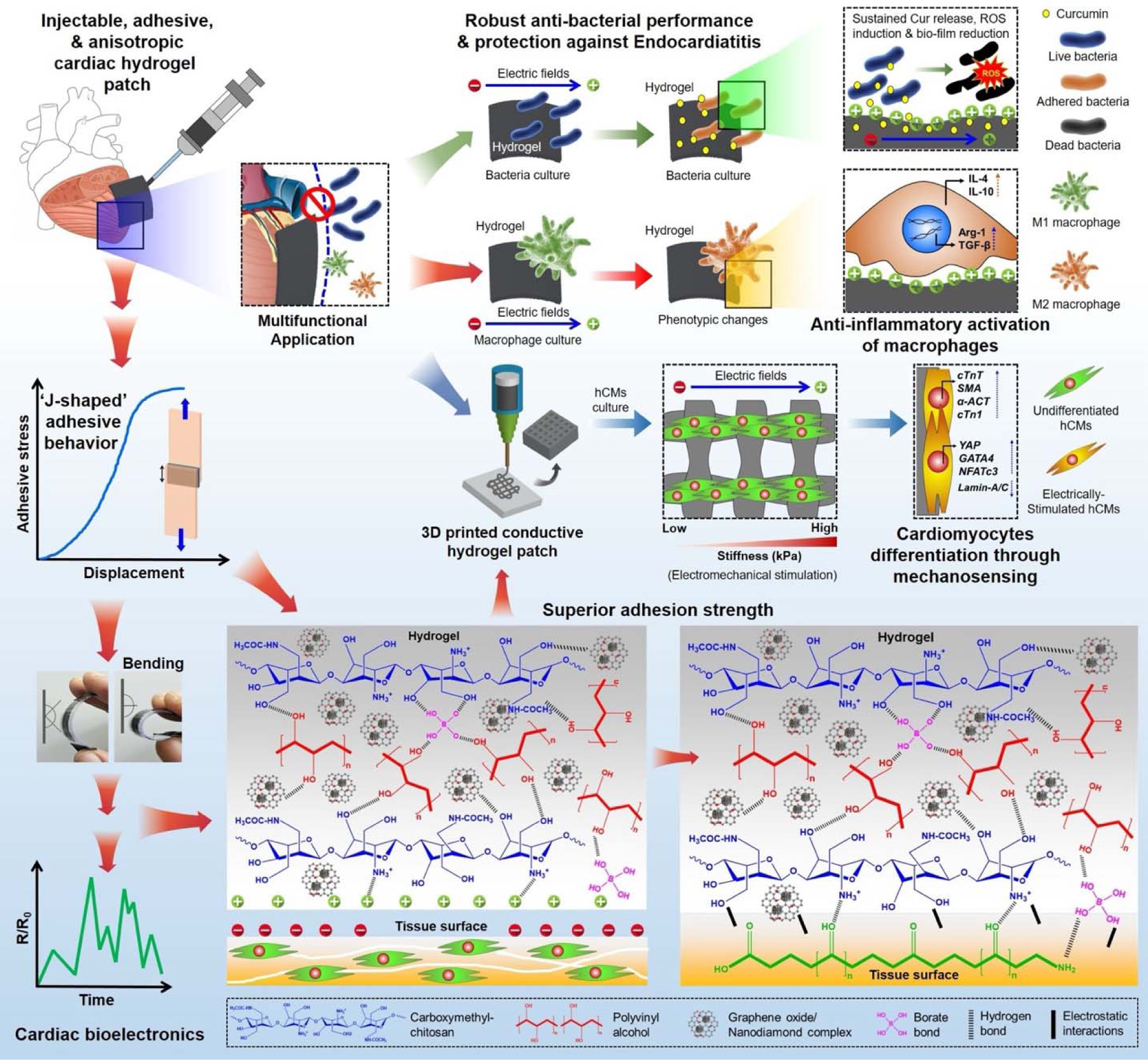
Schematic illustration of the 3D printable anisotropic and adhesive hydrogel patch with multifunctional abilities for regulating electromechanical and immunogenic cues of cardiac microenvironment.

## Notes

### Competing Interest Statement

The authors have declared no competing interest.

